# Axisymmetric diffusion kurtosis imaging with Rician bias correction: A simulation study

**DOI:** 10.1101/2022.03.15.484442

**Authors:** Jan Malte Oeschger, Karsten Tabelow, Siawoosh Mohammadi

## Abstract

**Purpose:** To compare the estimation accuracy of axisymmetric diffusion kurtosis imaging (DKI) and standard DKI in combination with Rician bias correction (RBC) under the influence of noise.

**Methods:** Axisymmetric DKI is more robust against noise-induced variation in the measured signal than standard DKI because of its reduced parameter space. However, its susceptibility to Rician noise bias at low signal-to-noise ratios (SNRs) is unknown. Here, we investigate two main questions: first, does Rician bias correction improve estimation accuracy of axisymmetric DKI?; second, is the estimation accuracy of axisymmetric DKI increased compared to standard DKI? Estimation accuracy was investigated on the five axisymmetric DKI tensor metrics (AxTM): the parallel and perpendicular diffusivity and kurtosis and the mean kurtosis, using a simulation study based on synthetic and in-vivo data.

**Results:** We found that RBC was most effective for increasing accuracy of the parallel AxTM in highly to moderately aligned white matter. For the perpendicular AxTM, axisymmetric DKI without RBC performed slightly better than with RBC. However, the combination of axisymmetric DKI with RBC was the overall best performing algorithm across all five AxTM and the axisymmetric DKI framework itself substantially improved accuracy in tissues with low fiber alignment.

**Conclusion:** The combination of axisymmetric DKI with RBC facilitates accurate DKI parameter estimation at unprecedented low SNRs (≈ 15), possibly making it a valuable tool for neuroscience and clinical research studies where scan time is a limited resource. The tools used in this paper are publicly available in the open-source ACID toolbox for SPM.

## 1 Introduction

Diffusion weighted MRI is an in-vivo imaging modality used in neuroscience and clinical research. It is sensitive to changes in nervous tissues that, e.g., go along with neurodegenerative diseases like epilepsy and multiple sclerosis (Deppe et al., 2008; Rovira et al., 2015). Diffusion MRI measures the net diffusion of nuclear spins of hydrogen nuclei in water molecules that are omnipresent in nervous tissue.

Diffusion of water molecules within the microstructural tissue landscape can be arbitrarily complex. A data-efficient method that captures both, standard Gaussian diffusion and more complex restricted diffusion processes (e.g., due to diffusion of water trapped in the cell-body of axons), is the recently introduced axisymmetric diffusion kurtosis imaging (DKI) framework (Hansen et al., 2016; Hansen and Jespersen, 2017). Its data-efficiency stems from requiring fewer diffusion weighted images than standard DKI, for the following reasons: while standard diffusion kurtosis imaging (DKI) is based on 22 parameters (Jensen et al., 2005), axisymmetric DKI has a reduced parameter space of 8 parameters due to imposing the assumption of axisymmetrically distributed axons. This is likely a reasonable assumption in major white matter fiber bundles (Hansen et al., 2016). Furthermore, this is expected to make axisymmetric DKI less susceptible to the noise induced variation of the acquired diffusion MRI signals.

Noise in MRI images introduces a random variation into the measured diffusion signals and a bias for the estimated DKI parameters when the signal-to-noise-ratio (SNR) is low. This bias is known as the “Rician noise bias” (Henkelman, 1985; Gudbjartsson and Patz, 1995; Sijbers et al., 1998) and becomes more severe, the lower the SNR is. Diffusion MRI is prone to a low SNR (Derek K. Jones, 2012) because it generates image contrast from additional spin dephasing associated with water mobility leading to a signal attenuation. DKI is even more susceptible to the Rician noise bias compared to conventional diffusion tensor imaging (DTI), since estimating the DKI parameters requires multiple diffusion shells including higher diffusion weighting, lowering the SNR. This increases the demand for effective Rician bias correction (RBC) schemes (Veraart et al., 2013a; Glenn et al., 2015) in DKI. Right now it is unclear whether fitting the axisymmetric DKI framework with its reduced parameter space (axisymmetric DKI has 8 parameters while standard DKI has 22) is better suited for parameter estimation from noisy diffusion MRI data than standard DKI. And it is furthermore unknown, how this could affect the susceptibility to the Rician noise bias.

The effect of the Rician noise bias on the fractional anisotropy (FA), mean diffusivity (MD), mean kurtosis 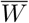, diffusion tensor and diffusion kurtosis tensor elements was shown to be mitigated by using RBC in standard DKI (Koay et al., 2009; Veraart et al., 2011, 2013a; André et al., 2014).

Of these parameters, only the mean kurtosis is part of the axisymmetric DKI tensor metrics (AxTM). The AxTM are the parallel and perpendicular diffusivities (*D*_∥_ and *D* _⊥_) and the mean, parallel, and perpendicular kurtosis (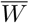, *W*_∥_ and *W*_⊥_). Here parallel and perpendicular are in reference to the axis of symmetry. The axisymmetric DKI framework contains three additional parameters, the two angles of the unit vector pointing along the axis of symmetry, and the non diffusion-weighted signal (*b* = 0).

The AxTM can be estimated based on standard DKI where they are computed as aggregates from its 22 parameters or directly with axisymmetric DKI (see Table 2.1). The AxTM are of particular interest for neuroscience and clinical research (Coutu et al., 2014; Genç et al., 2018; Donat et al., 2021) because they are invariant against coordinate transformations and describe free and restricted diffusion within nervous tissue. Furthermore, the AxTM can be directly related to the tissue microstructure. The latter relation is established via their mathematical relation to the five microstructure parameters of the biophysical standard model (Novikov et al., 2018; Jespersen et al., 2018): axon water fraction, axon dispersion, and three diffusivities associated with the intra- and extra-axonal space.

**Table 2.1:**
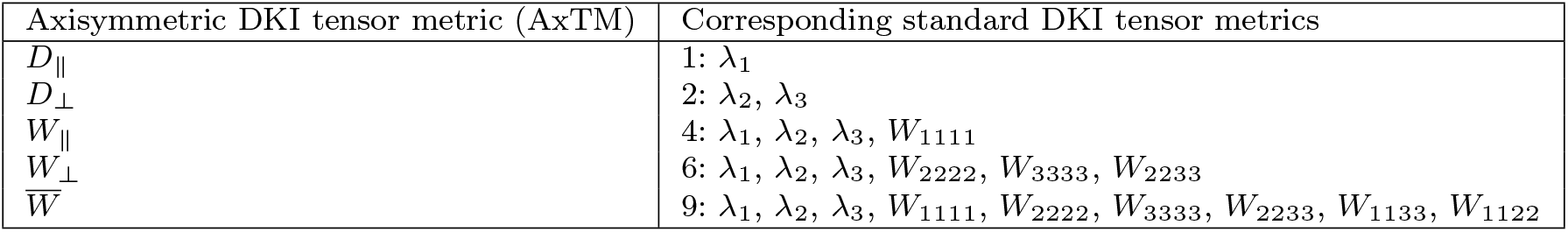
AxTM and standard DKI tensor metrics with which they are calculated. The numbers show how many standard DKI tensor metrics are needed to compute the AxTM. λ refers to the eigenvalues of the diffusion tensor, *W* refers to the components of the kurtosis tensor.

| Axisymmetric DKI tensor metric (AxTM) | Corresponding standard DKI tensor metrics |
| --- | --- |
| $D_{\parallel}$ | 1: $\lambda_1$ |
| $D_{\perp}$ | 2: $\lambda_2, \lambda_3$ |
| $W_{\parallel}$ | 4: $\lambda_1, \lambda_2, \lambda_3, W_{1111}$ |
| $W_{\perp}$ | 6: $\lambda_1, \lambda_2, \lambda_3, W_{2222}, W_{3333}, W_{2233}$ |
| $\bar{W}$ | 9: $\lambda_1, \lambda_2, \lambda_3, W_{1111}, W_{2222}, W_{3333}, W_{2233}, W_{1133}, W_{1122}$ |

Furthermore, it was shown empirically (Veraart et al., 2011) that RBC will impact the estimation of the parallel and perpendicular AxTM differently. It was speculated that the parallel and perpendicular AxTM are associated with different levels of water mobility and consequently different levels of SNR. Another open question is the influence of fiber alignment on the effectiveness of RBC. It can be expected that the degree of fiber alignment within a white matter MRI voxel affects water mobility and thereby also the efficiency of RBC.

In this work two main questions are investigated: First, and motivated by the improved parameter estimation accuracy when applying RBC in standard DKI, we investigate whether RBC also increases the estimation accuracy of axisymmetric DKI. Second, we investigate whether the estimation accuracy is improved by using axisymmetric DKI as compared to standard DKI. Moreover, we investigate whether the performance of RBC depends on tissue fibre alignment in a voxel and investigate differences in effectiveness for the parallel and perpendicular AxTM. To study these questions, we simulated two datasets: the “`synthetic dataset`” is based on three sets of synthetic AxTM describing tissues with varying degrees of fiber alignment which allows us to assess AxTM estimation accuracy as a function of fiber alignment; the “`in-vivo like dataset`” is based on in-vivo measurements of white matter tissue fiber tracts with a high to moderate fiber alignment which allows us to study AxTM estimation accuracy under realistic, in-vivo like conditions. In both studies, axisymmetric DKI and standard DKI (with and without RBC) were used to obtain estimates of the five AxTM.

## 2 Methods

### 2.1 Standard DKI signal representation

For a given diffusion weighting *b* and diffusion gradient 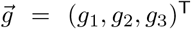, the noise-free DKI signal can be represented as Jensen and Helpern (2003); Jensen et al. (2005):

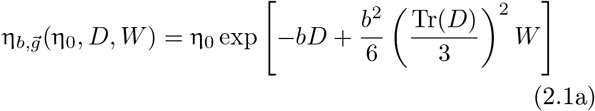

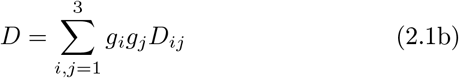

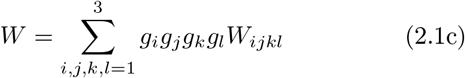

where *D*_*ij*_ are the diffusion tensor entries, *W*_*ijkl*_ are the kurtosis tensor entries and η_0_ is the non-diffusion-weighted signal 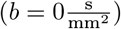.

### 2.2 Axisymmetric DKI

Axisymmetric DKI (Hansen et al., 2016) assumes symmetric diffusion around an axis of symmetry 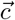 inside an imaging voxel. Mathematically, this assumption leads to axisymmetric diffusion and kurtosis tensors with a drastically reduced number of independent tensor parameters compared to standard DKI (from 15 to 3 parameters for the kurtosis tensor and from 6 to 2 parameters for the diffusion tensor). The symmetry assumptions are likely to be a reasonable approximation to diffusion in major white matter fiber bundles (Hansen et al., 2016) due to their structural organization.

With the axis of symmetry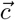 parameterized by the inclination *θ* and azimuth 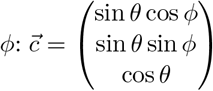, the diffusion and kurtosis tensors can be determined according to Hansen et al. (2016):

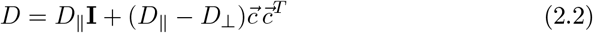

and

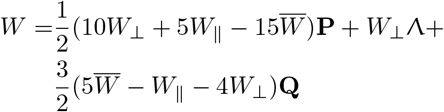

where 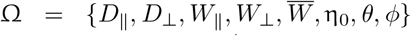 are the 8 framework’s parameters (η_0_ is the non diffusion-weighted signal) and 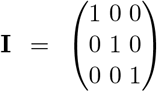 is the identity matrix. The tensors **P**, Λ and **Q** can be computed with the Kronecker delta δ_*xy*_ and the components of the axis of symmetry *c*_*x*_ (*x, y* ∈ 1, 2, 3) as: **P**_*ijkl*_ = *c*_*i*_*c*_*j*_*c*_*k*_*c*_*l*_, 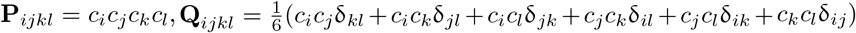 and 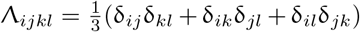 (Hansen et al., 2016). The according noise-free signal 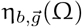 can then be computed based upon the axisymmetric tensors (Hansen et al., 2017):

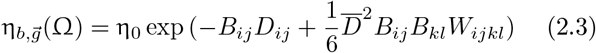

where

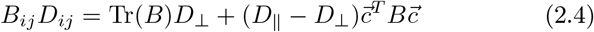

and

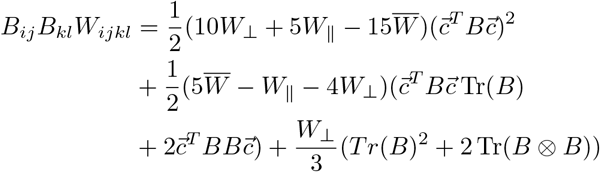

with 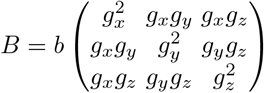.

The AxTM can be computed from the standard DKI tensor metrics assuming axial-symmetry. For example, 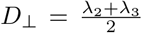 where λ refers to the eigenvalues of the diffusion tensor. Formulas for the computation of the other AxTM can be found in Tabesh et al. (2011), Table 2.1 shows the AxTM and the standard DKI tensor metrics needed to compute them. Figure 2.1 shows the five AxTM obtained with the axisymmetric DKI fit without RBC, available in the open source ACID toolbox for SPM.

**Fig. 2.1:**
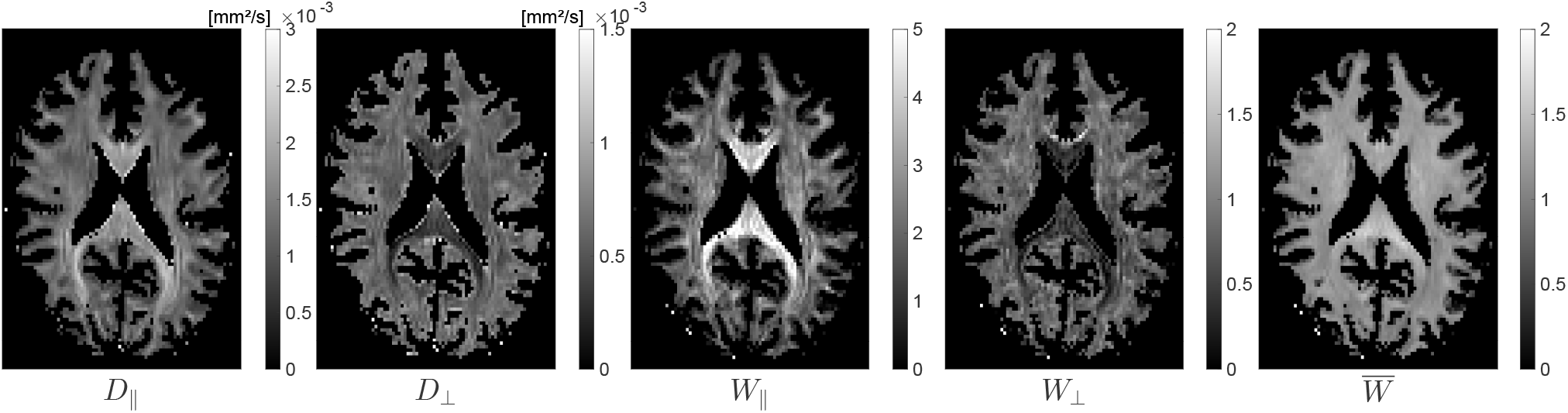
AxTM results in white matter. The AxTM are the parallel and perpendicular diffusivity and kurtosis and the mean kurtosis. The shown maps were obtained with the axisymmetric DKI fit available in the open source ACID toolbox for SPM that was used in this work. The AxTM were estimated from the in-vivo measurement used for the `in-vivo like dataset` (Figure 2.3).

### 2.3 Parameter estimation and the Rician noise bias

Standard DKI or axisymmetric DKI parameter estimation would typically be done using the acquired magnitude signals 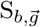 and Eq. (2.1a) or Equation (2.3) in the least-squares approach (Veraart et al., 2013b; Tabesh et al., 2011):

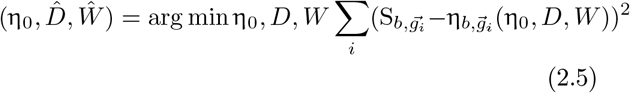

However, this least-squares approach is built on the assumption of Gaussian distributed noise in 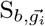 which is not true in reality. 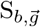 is a magnitude signal computed from the noise contaminated k-space data described by a complex Gaussian (standard deviation σ) as the sum of squares of the measured signal intensity (Constantinides et al., 1997) from the receiver coil after it was Fourier transformed into real space. Computing the sum of squares rectifies the composite magnitude signal and leads to Rician distributed noise for one receiver coil (L= 1). Therefore, assuming Gaussian noise in MRI magnitude signals leads to a bias that propagates into the estimated parameters which is referred to as the “Rician noise bias”. Eq.(2.5) is therefore biased.

More generally, if one assumes uncorrelated noise and statistically independent receiver coils with an equivalent noise variance (Aja-Fernández and Tristán-Vega, 2012), the resulting probability density function of the noisy magnitude data is given by a non-central χ-distribution (Constantinides et al., 1997), where 2*L* is the number of degrees of freedom of the distribution. *L* = 1 results in the Rician distribution (Rice, 1944; Gudbjartsson and Patz, 1995).

The severity of the Rician noise bias depends on the SNR (Polzehl and Tabelow, 2016) because the sum of squares rectifies the composite magnitude signal: the lower the SNR, the larger the bias. For RBC, we rely on an approach outlined in Polzehl and Tabelow (2016) that uses the expectation value 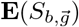 of the composite magnitude signal. The probability density function of 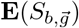 is a non-central χ distribution and given by (Polzehl and Tabelow, 2016):

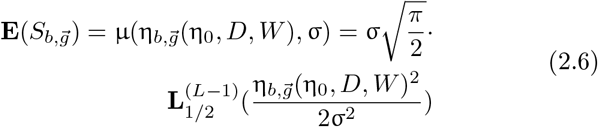

where 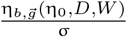 is the SNR and 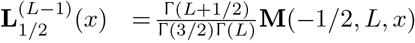 is the generalized Laguerre polynomial which can be expressed using a confluent hypergeometric function **M** and the Gamma function Γ. Only for simplicity of notation, in the text we neglect any possible dependence of *σ* on b, 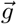 or location, the employed RBC algorithm used the same *σ* in every image voxel. The SNR dependent expectation value Eq. (2.6) differs from the noise-free signal, 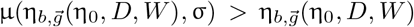 with the difference decreasing with increasing SNR. Following Polzehl and Tabelow (2016), we implemented a time-efficient fitting algorithm that, unlike Eq. (2.5), accounts for Rician noise in magnitude MRI data by solving the optimization problem:

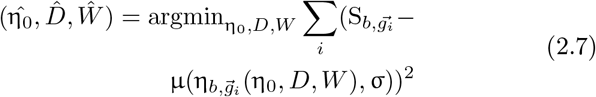

Estimating parameters this way is referred to as “quasi-likelihood” estimation and is denoted as “RBC ON” in this paper. It was shown, that parameter estimation using the non-central χ noise statistic in a quasi-likelihood framework yields asymptotically unbiased parameter estimates (Bunke and Schmidt, 1980; Polzehl and Tabelow, 2016).

Rician bias corrected, standard DKI or axisymmetric DKI parameter estimation can be done by using Eq. (2.1a) or Eq. (2.3) to compute the noise-free signal predictions 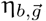, then using Eq. (2.6) to compute 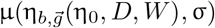 and finally minimize Eq. (2.7) to estimate the framework parameters (η_0_, *D, W*) for standard DKI or Ω for axisymmetric DKI.

In reality, noise correlations between receiver coils occur and are non-negligible, especially for a higher number of receiver coils (32 or 64). This affects the degrees of freedom of the underlying noise statistic. However, the non-central χ distribution can still be used as a good approximation, if an effective number of coils *L*_eff_ and noise variance 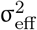 are used (Aja-Fernández and Tristán-Vega, 2012) for which *L ≥ L*_eff_ and 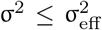 can be shown. Similarly, the generalized autocalibrating partially parallel acquisition (GRAPPA) scheme can be accounted for by specifying an effective number of coils *L*_eff_, while *L* = 1 for sensitivity encoding (SENSE) (Aja-Fernández and Tristán-Vega, 2012).

### 2.4 Parameter estimation with the Gauss-Newton algorithm

To minimize Eq. (2.5) or Eq. (2.7) time-efficiently, we have implemented a Gauss-Newton minimization algorithm (Modersitzki, 2009) in Matlab for slice-wise and parallelizable parameter estimation on MR-images instead of using standard Matlab optimization functions. The used tools are freely available online within the ACID toolbox (http://www.diffusiontools.com/) for SPM. Slice-wise fitting refers to fitting all voxels of an image-slice at the same time which improves run-time. The implemented algorithm is highly adaptable and can fit any signal model (especially non-linear models). Gauss Newton parameter estimation approximates the search direction in parameter space based on the Jacobian and is sensitive to the initial guess. For the initial guess of the axisymmetric DKI fit implementation, we used code from the repository of Sune Nørhøj Jespersen: https://github.com/sunenj/Fast-diffusion-kurtosis-imaging-DKI (Hansen et al., 2016).

### 2.5 Simulation study: Datasets and overview

We assessed estimation accuracy of the five AxTM as a function of the SNR in a simulation study with two types of datasets. One dataset consisted of three synthetic voxels with varying fiber alignment (defined in Coelho et al. (2019)). This dataset is refereed to as “`synthetic dataset`” because it was derived in the context of another study (Coelho et al., 2019) by random sampling of the parameter space of biophysical parameters and consequent derivation of the corresponding AxTM. The other dataset consisted of twelve major white matter fiber tract voxels from an in-vivo brain measurement, this dataset is refereed to as “`in-vivo like dataset`”. Details on both datasets are listed below and in Figure 2.2. For both datasets, magnitude diffusion MRI data were simulated for varying SNRs and fitted with standard DKI and axisymmetric DKI, with and without RBC (as described in Section 2.3) to obtain estimates of the five AxTM. Accuracy of the obtained AxTM estimates were evaluated as the mean absolute percentage error (MAPE):
1

**Fig. 2.2:**
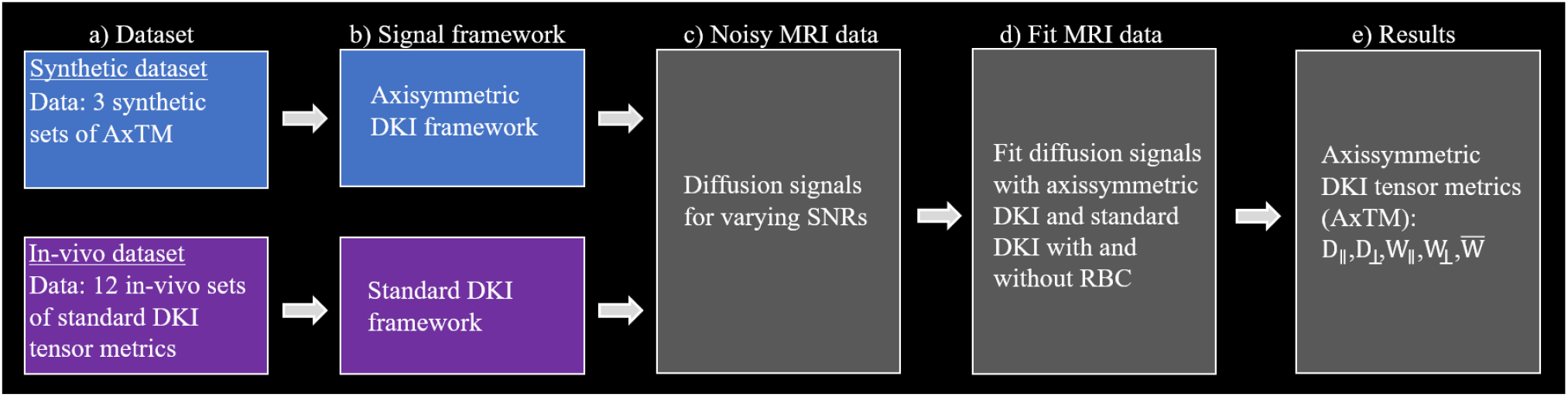
Scheme of the simulation study. Simulations were performed in 5 steps: a) choice of datasets, b) signal framework used for simulation, c) diffusion signal simulation and contamination with noise, d) parameter estimation and e) results. Note that both simulation studies only differed in a) and b) but were identical in the following procedures. a) the `synthetic dataset` consisted of 3×5 sets of AxTM (Table 8.3) while the `in-vivo like dataset` consisted of 12×22 standard DKI tensor metrics (Table 8.1 in the Supplementary material). b) The DKI signal framework used for diffusion signal simulation. c) Diffusion signal data were contaminated with 2500 Rician noise samples for each SNR=[1, 2, 3…100]. d) Simulated diffusion data were fitted with axisymmetric DKI and standard DKI with and without RBC in both simulation studies for each of the 2500 noise samples. e) The axisymmetric DKI tensor metrics (AxTM): *D*_∥_, *D*_⊥_, *W*_∥_, *W*_⊥_ and 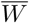 were calculated for standard DKI data (for axisymmetric DKI they were directly estimated), averaged across the 2500 noise samples per SNR and finally compared to the ground truth.

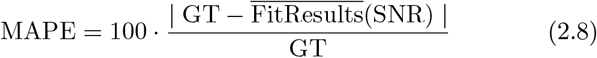

Here GT reefers to the ground truth and 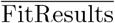 reefers to the average of the fit results over the noise samples. We evaluated the accuracy of the AxTM estimates for each estimation method by looking for the SNR after which the MAPE was smaller 5%. The 5% threshold was considered an acceptable error in a trade-off between estimation accuracy and SNR requirement. The different setup of both simulation studies enables an isolated investigation of the effectiveness and tissue dependence of the RBC and to test the fitting methods in an `in-vivo like dataset`. As a summary to compare each method, we looked at the maximum SNR needed across the five AxTM for which MAPE consistently *<* 5% for all AxTM (“Maximum” column in Figure 3.2).

#### Datasets

The `synthetic dataset` consisted of three synthetic sets of AxTM (from Coelho et al. (2019)) describing three voxels with varying fiber alignment, one with fibers with low alignment (“LA”, FA=0.067), one with fibers with moderate alignment (“MA”, FA=0.24) and one with highly aligned fibers (“HA”, FA=0.86). The AxTM of the three synthetic voxels are summarized in Table 8.3 (“Supplementary material”). Figure 2.4 shows two areas of typical brain regions in a map of the mean kurtosis 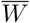 where LA and HA voxels can be found and the corresponding idealized fiber stick model.

The `in-vivo like dataset` consists of twelve voxels extracted from four major white matter tracts (three voxels from each of the four fiber tracts, see Figure 2.3) from an in-vivo brain measurement (SNR=23.4) of a healthy volunteer. The twelve voxels were extracted from the in-vivo measurement by fitting the standard DKI framework in 12 voxels to get the corresponding 22 standard DKI tensor metrics, the derived data are therefore refereed to as “in-vivo like”. Three voxels with HA to MA (defined through their fractional anisotropy (FA)) were extracted from these four major white matter fiber tracts based upon the Jülich fiber atlas: the callosum body (`cb`), the corticospinal tract (`ct`), the optic radiation (`or`) and the superior longitudinal fasciculus (`slf`), see Figure 2.3. The sets of the 12 in-vivo like standard DKI tensor metrics are documented in the Supplementary material in Table 8.1, the derived AxTM are found in Table 8.2.

**Fig. 2.3:**
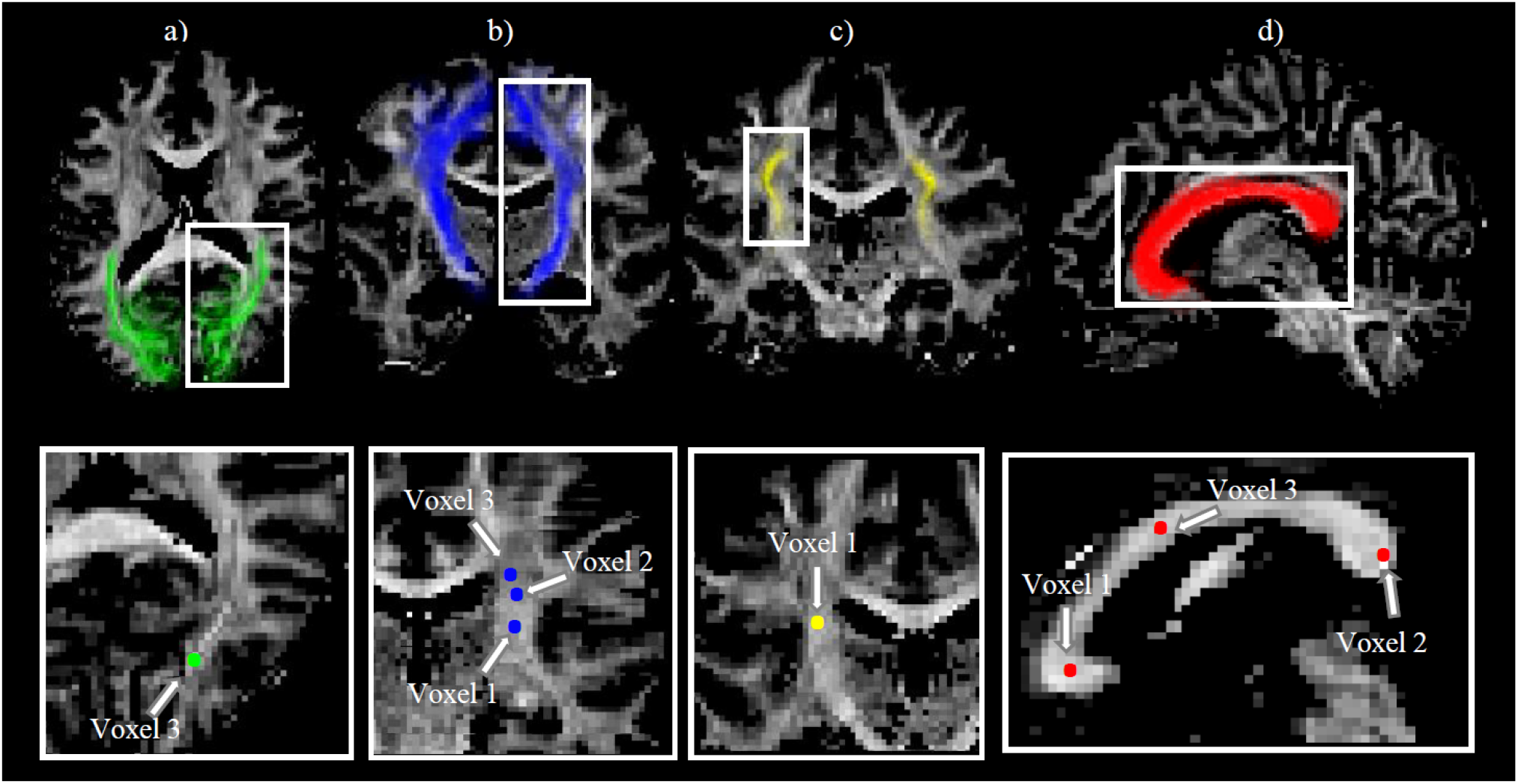
Selection of voxels of the `in-vivo like dataset`: The four white matter fiber pathways within which voxels were selected and used as a basis for the `in-vivo like dataset`. Top: a) Optic radiation (or), b) cortico spinal tract (ct), c) superior longitudinal fasciculus (slf) and d) callosum body (cb) in a fractional anisotropy (FA) map of a healthy human brain. The fiber pathways were identified with the coregistered Jülich fiber atlas (Eickhoff et al., 2005). Bottom: Voxels in the fiber pathways used for the `in-vivo like dataset`. In each fiber pathway, three voxels were chosen for the `in-vivo like dataset` (for slf an or only one is shown here because the chosen voxels were not in the same slice).

#### Signal framework used for simulation

The three synthetic voxels of AxTM were simulated with the axisymmetric DKI framework to first obtain noise-free diffusion MRI magnitude signals. The twelve in-vivo like voxels were simulated with the standard DKI framework to first obtain noise-free diffusion MRI magnitude signals.

#### Contamination with noise

For both the synthetic and the `in-vivo like dataset`, the noise-free diffusion MRI magnitude signals were contaminated with noise for SNRs [1, 2, 3…100].

#### Estimating the five AxTM

Both, the simulated signals from the `synthetic` and the `in-vivo like dataset` were fitted with axisymmetric DKI and standard DKI, with and without RBC (section 2.3) to obtain estimates of the AxTM whose accuracy could then be investigated as a function of SNR.

### 2.6 In-vivo data acquistion and simulated sequence

The DWI sequence used to acquire the `in-vivo like dataset` was a mono-polar single-shot spin-echo EPI scheme, consisting of 16 non-diffusion-weighted images (*b* = 0 image). The diffusion weighted images were acquired at three b values 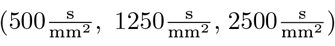, sampled for 60 unique diffusion-gradient directions for the 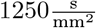 and 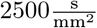 shells and 30 unique directions for the 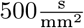 shell. The entire protocol was repeated with reversed phase encoding directions (“blip-up”, “blip-down” correction) to correct for susceptibility-related distortions so that in total 166*·* 2 images were acquired. Other acquisition parameters were: an isotropic voxel size of (1.6mm^3^), FoV of 240×230×154mm^3^, TE = 73ms, TE = 5300ms and 7/8 partial Fourier imaging. Signal simulation in our simulation study was done with only one *b* = 0 signal, so that the simulated sequence consisted of 151 signals per noise realization.

### 2.7 Simulation studies: Details

We simulated 100 SNRs: SNR = [1, 2, 3, …100]. Noise was added according to S_cont_ = |S_noise−free_ + α + β*i*|, where α, β ∈ *𝒩* (0, σ) are drawn from a zero mean Gaussian with standard deviation σ, yielding different 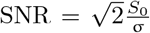 (for one receiver coil) for a given *S*_0_ = 1. For every SNR, 2500 noise samples were realized, i.e., 2500 *·* 151 pairs (α, β) were drawn and 2500 *·* 151 S_cont._ were calculated per SNR for every simulated voxel. These diffusion MRI magnitude signals were then fitted with the four proposed methods (section 2.3). For each of the 2500 noise samples per SNR, 2500 parameter estimates of 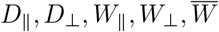 were obtained and averaged to find the SNR above which the average over these 2500 noise samples had a MAPE *<* 5% (`synthetic datset`). For the `in-vivo datset` this MAPE was averaged per SNR across the 12 simulated voxels and the SNR above which this averaged MAPE *<* 5% is reported.

For simulation of the three synthetic voxels, the axis of symmetry 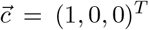 was fixed throughout the study. For data fitting, the two angles *θ* and *ϕ* that define the axis of symmetry within the axisymmetric DKI framework were variable but constrained to *θ, ϕ* ∈ [−2*π*, 2*π*] which improved convergence of the fitting algorithm. Data were simulated according to the simulation scheme described in Section 2.6.

### 2.8 Diffusion signal profiles influenced by fiber alignment

To further elucidate differences between tissues with different levels of fiber alignment, angular signal profiles under the influence of noise were studied for the three voxels of the `synthetic dataset`. Noise-free and noise-contaminated signals have been simulated (SNR = 20). The simulated signal’s mean and standard deviation could then be plotted as a function of angle *ψ* (in degree) between diffusion gradient 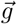 and axis of symmetry 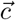. For a graphical representation of angle *ψ*, see Figure 2.4.

**Fig. 2.4:**
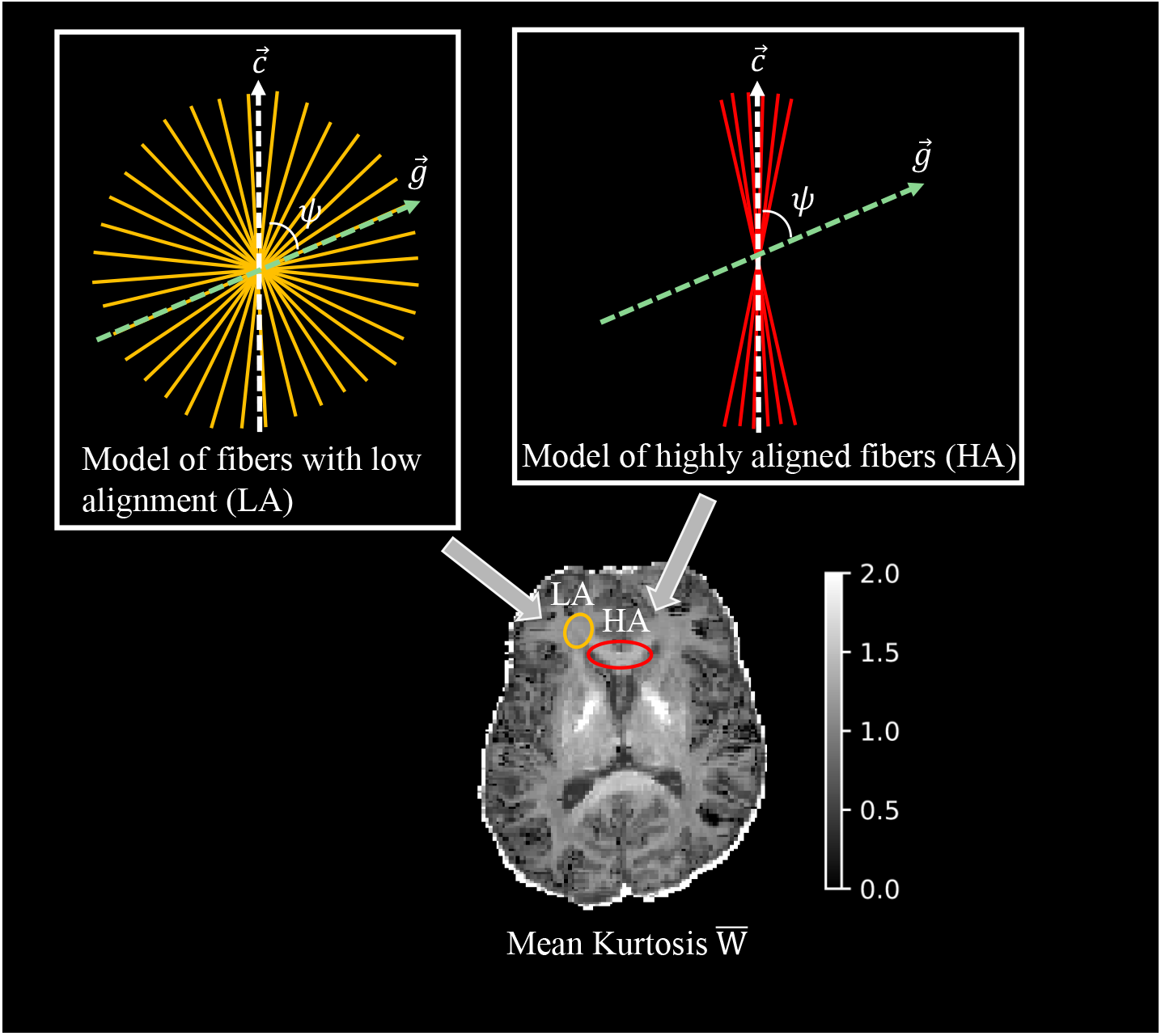
Model of fiber alignment in characteristic areas of the brain. Left: in-vivo map of the mean kurtosis 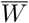 with a typical area where fibers with low alignment (LA) are found and a typical area where highly aligned fibers (HA) are found. Right: the corresponding golden and red sticks depict an idealized model of the underlying fiber arrangement, the white dashed line indicates the axis of symmetry 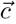, the green dashed line indicates the diffusion gradient direction 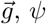 is the angle between 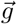 and 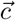.

## 3 Results

First, the results of the diffusion signal profiles in voxels with different levels of fiber alignments (Section 3.1) are shown because these not only explain the results obtained in different tissues but also help to understand the difference between estimating the parallel or the perpendicular AxTM. After that, our main findings are stated and the corresponding results are reported (Section 3.2).

### 3.1 Diffusion signal profiles influenced by fiber alignment

Each of the simulated voxels of the `synthetic dataset` shows a characteristic, *ψ* dependent shape, see Figure 3.1. For smaller angles *ψ* between *ψ* = 20^*°*^ and *ψ* = 0^*°*^, the simulated signals of white matter with HA are strongly diffusion weighted and are close or below the noise floor (SNR = 1) where the signal and noise strength are equal, see Figure 3.1a. In the simulated voxel of MA, the noise floor is already reached for angles *ψ* ≈ 50^*°*^, see Figure 3.1b. In the simulated voxel of white matter with low fiber alignment the noise floor is never reached and the simulation shows a seemingly constant, *ψ* independent signal form, see Figure 3.1c. In summary, the signal in HA to MA decays along the direction of symmetry, whereas there is almost no decay in LA.

**Fig. 3.1:**
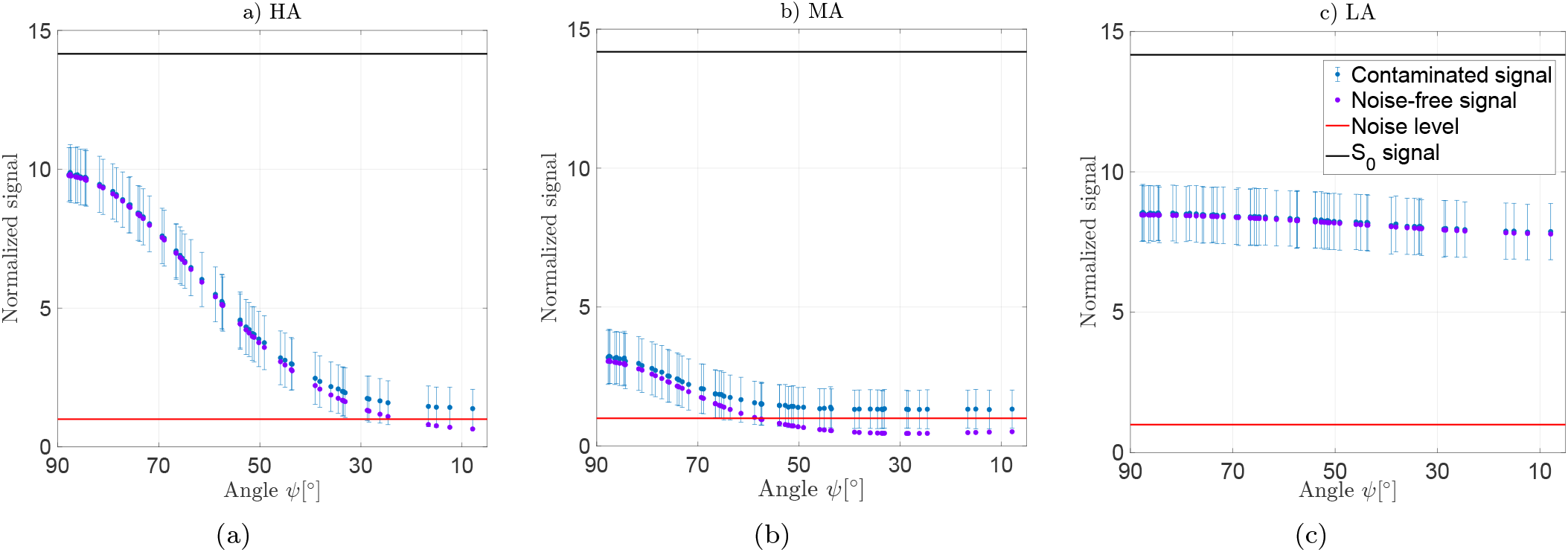
Simulated signal decay along the axis of symmetry at SNR = 20. Signal decay is shown for the `synthetic dataset` consisting of a) highly aligned fibers (HA), b) fibers with moderate alignment (MA) and c) fibers with low alignment (LA) as a function of angle *ψ* between the diffusion gradient 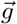 and the axis of symmetry 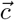, see Section 2.8. The “Contaminated signal” shows the mean and standard deviation over 2500 noise-samples. All signals, including the noise-free ones, were normalized to the noise strength, the plots always show 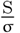. The SNR is calculated for the signal at 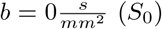 according to: 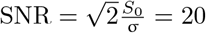 which corresponds to 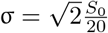. All three plots show the signals for the highest b-shell 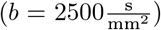. For a graphical representation of angle *ψ* see Figure 2.4.

### 3.2 Results of simulation study

Figure 3.2 shows the SNRs required to accurately estimate the AxTM (MAPE*<* 5%, Equation (2.8)) in the `synthetic` and `in-vivo like dataset`. Shown are the results for axisymmetric DKI and standard DKI with (hatched) or without RBC.

**Fig. 3.2:**
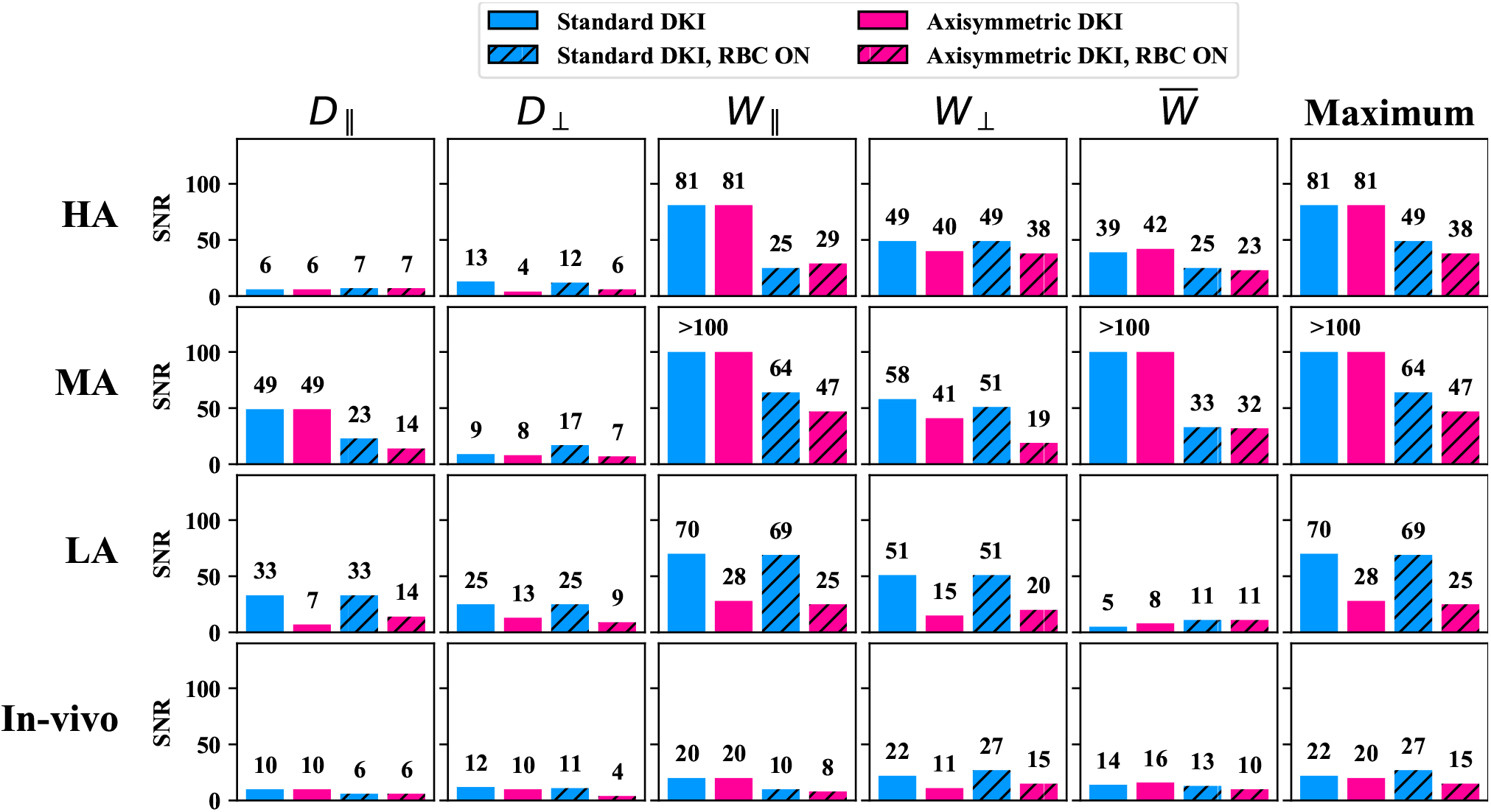
Signal-to-noise ratio (SNR) above which the mean absolute percentage error (MAPE, Equation (2.8)) *<* 5% for the `synthetic dataset` with high, medium and low fiber alignment (“HA”, “MA”, “LA”) and the `in-vivo like dataset`. For the `in-vivo like dataset`, the MAPE was averaged across the 12 simulated voxels and the SNR above which this average MAPE *<* 5% is shown. The standard deviation across the in-vivo dataset voxels is not shown here because the values were between ≈ 0.5 and ≈ 6 with an average of ≈ 2.5 and thus to small to display. The number above the barplots indicates the barplot’s height. Blue encodes standard DKI, red encodes axisymmetric DKI, the hatched barplots show the results if RBC is used. “Maximum” shows the maximum SNR needed to achieve MAPE *<* 5% across all five AxTM.

#### RBC most effective in highly aligned fibers and parallel diffusion

RBC was most effective for the parallel parameters *D*_∥_ and *W*_∥_ in HA to MA (for both the synthetic and `in-vivo like dataset`), see Figure 3.2. For example, achieving MAPE <5% for *W*_∥_ in the synthetic HA voxel could be reduced from SNR= 81 to SNR=25 (standard DKI) or SNR=29 (axisymmetric DKI). In the in-vivo like voxels, the SNR requirements for achieving MAPE <5% for *W*_∥_ could be reduced from 20 to 10 (standard DKI) or to 8 (axisymmetric DKI). Estimation of 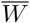 was significantly improved by RBC in the synthetic HA and MA voxels but only slightly in the `in-vivo like datasets`.

#### Superiority of axisymmetric DKI in fibers with low alignment where RBC is ineffective

In the synthetic LA voxel, estimation of *D*_∥_, *D*_⊥_, *W*_∥_ and *W*_⊥_ were substantially improved by using the axisymmetric DKI framework instead of standard DKI. E.g., it only required an SNR= 15 (axisymmetric DKI) instead of SNR= 51 (standard DKI) to achieve MAPE <5% for *W*_⊥_. For 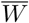, axisymmetric DKI performed slightly worse than standard DKI (SNR= 8 instead of SNR= 5). Interestingly, RBC did not influence the fitting results much in this fiber alignment configuration but even worsened them in some cases (e.g. *D*_∥_), see Figure 3.2.

#### Axisymmetric DKI improves estimation of perpendicular parameters

Estimation of *D*_⊥_ and *W*_⊥_ could also be improved (MAPE <5% reached for lower SNRs) by using the axisymmetric DKI framework in HA to MA of the `synthetic` and `in-vivo like dataset` (e.g., for HA, *D*_⊥_: SNR= 4 instead of SNR= 13; *W*_⊥_: SNR= 40 instead of SNR= 49).

#### RBC can also worsen accuracy

Interestingly, there were also few scenarios in which RBC increased the SNR requirements. As described above this was observed for *D*_∥_, *W*_⊥_ and 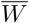 in the LA voxel and, outside the LA voxel, predominantly for the perpendicular parameters, e.g., from SNR= 11 to SNR= 15 for the axisymmetric DKI fit of *W*_⊥_ in the `in-vivo like dataset`.

## 4 Discussion

### 4.1 Summary of main findings

Overall, we found, that the combination of axisymmetric DKI with Rician bias correction (RBC) was the best option for estimating all five axisymmetric DKI tensor metrics (AxTM), see “Maximum” column of Figure 3.2. This combination achieved a mean absolute percentage error (MAPE, Equation (2.8)) *<* 5%) on our simulated `in-vivo like data` if the signal-to-noise ratio (SNR) *≥* 15, making this combination a possibly valuable tool in neuroscience and clinical research studies. Specifically, we found that RBC is highly effective for increasing estimation accuracy of the AxTM associated with diffusion parallel to the main fiber orientation, i.e., parallel diffusivity and kurtosis, in white matter with HA. In contrast, it fails in improving estimation accuracy in parameters perpendicular to the main fiber orientation, i.e., perpendicular diffusivity and kurtosis, or if fiber alignment is too low. For the latter scenarios, axisymmetric DKI is more effective than standard DKI.

### 4.2 Rician noise bias and its correction: Effectiveness for different DKI parameters and levels of fiber alignment within white matter

The effectiveness of RBC correlated with the severity of the Rician noise bias influenced by either the level of fiber alignment or direction of the AxTM which indicates that the proposed method for RBC indeed mitigates Rician noise bias. Severity of the Rician noise bias is inversely proportional to the SNR and hence depends on a variety of parameters. One of these parameters is the level of water mobility which tunes the diffusivity and thereby the diffusion weighted signal. Water mobility is influenced by a combination of the level of fiber alignment and orientation of the AxTM relative to the axis of symmetry 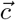. Figure 3.1 shows simulated signals in three tissue types with varying degrees of fiber alignment as a function of angle *ψ* between 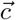 and diffusion gradient 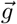. In HA for example, the diffusion signal is heavily diffusion weighted if measured along the main fiber orientation (small angle *ψ*) and the Rician noise bias in these signals therefore strongest, see Figure 3.1a. Since the parallel AxTM (*D*_∥_ and *W*_∥_) predominantly depend on these signals, it can be expected that they, too, are more heavily biased in a high fiber alignment setting. Accordingly, we found that Rician bias correction (RBC) turned out to be particularly important for the AxTM associated with parallel diffusion (*D*_∥_ and *W*_∥_) in highly aligned white matter (`synthetic dataset` and `in-vivo like dataset`). On the other hand, diffusion perpendicular to 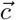 will be more restricted than the parallel diffusion and the SNR therefore higher. *D*_⊥_ and *W*_⊥_ should therefore be less affected by the Rician noise bias. Indeed, we found that RBC was not as effective for the perpendicular parameters *D*_⊥_ and *W*_⊥_ in white matter with HA as for the parallel parameters. An apparent contradiction to this argument was found for *W*_⊥_ in the synthetic MA dataset. This contradictory finding, however, can be explained by the relatively high perpendicular diffusivity of the MA voxel (Table 8.3) causing a strong diffusion weighting and therefore smaller perpendicular signal in this case (Figure 3.1b).

RBC was furthermore ineffective in tissues with LA (LA voxel of the `synthetic dataset` has an FA= 0.067), see Figure 3.2. Here, the signal change as a function of angle between 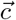 and 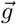 is smaller than the variation introduced by noise and the signals almost seem independent of the diffusion gradient direction. This is because these tissues do not posses a clearly distinguishable axis along which water mobility is significantly heightened compared to other directions, see Figure 3.1c. For the same SNR (in reference to the *S*_0_ signal), the Rician noise bias in such tissues is therefore less severe, compared to, e.g., signals in HA acquired for diffusion gradients parallel to the axis of symmetry. Accordingly, we found that RBC had little to no effect on parameter estimation in white matter with low fiber alignment. Furthermore, the original signal shape may be lost in the noise pattern which is why RBC could even increase the bias instead of reducing it because it acts on a falsely modeled signal form.

### 4.3 Advantage of the axisymmetric DKI framework

In the introduction, we hypothesized that a reduction of the parameter space could make axisymmetric DKI more robust against the Rician noise bias. We observed that axisymmetric DKI predominantly improved accuracy of parameter estimation of the perpendicular AxTM *D*_⊥_ and *W*_⊥_ in both the `synthetic dataset` (HA and MA voxels) and the `in-vivo like dataset`, i.e., MAPE *<* 5% was achieved for lower SNRs.

The increased accuracy for *W*_⊥_ can be understood with Table 2.1. In standard DKI, 6 of the 22 framework parameters are needed to compute *W*_⊥_, including the highly diffusion weighted and therefore highly Rician bias affected λ_1_. Axisymmetric DKI, on the other hand, always estimates *W*_⊥_ directly and therefore is getting rid of the highly Rician bias affected *λ*_1_ which mitigates the effects of Rician noise bias. Furthermore, the perpendicular AxTM are predominantly estimated from the “perpendicular signals” (see Section 4.2) which are less diffusion weighted in general. This additionally reduced effects of the Rician noise bias.

Another observation was that, accuracy of parameter estimation with axisymmetric DKI was substantially better in tissues with low fiber alignment (Figure 3.2) compared to standard DKI. We observed, e.g., an SNR reduction of up to 70% when using axisymmetric DKI instead of standard DKI for estimation of *W*_⊥_ in the LA voxel, see Figure 3.2. This could be due to the seemingly constant and high signal, independent of the diffusion gradient direction, in the synthetic LA dataset. Since the variation in the almost constant diffusion signal is dominated by noise, the complex 22 parametric standard DKI framework is more likely to overfit the data than the 8 parametric axisymmetric DKI framework, particularly in case of lower SNRs where noise has a greater impact. This could be the reason for the clear advantage of axisymmetric DKI over standard DKI in this fiber configuration.

### 4.4 Considerations

#### Possible circularity of simulation study

Since the simulation studies were either based on the axisymmetric DKI (`synthetic dataset`) or the standard DKI framework (`in-vivo like dataset`), one might argue that the simulations will favor their respective signal frameworks. Here we re-discuss our signal-framework comparisons in the light of this potential circularity: We observed that using axisymmetric DKI was advantageous over standard DKI for the perpendicular AxTM. The same trend was observed in the synthetic LA dataset. Since the improvement of axisymmetric DKI over standard DKI for the perpendicular AxTM was observed across both simulations, the observation cannot be explained by a circularity argument and we believe that it is a genuine advantage of axisymmetric DKI. The synthetic LA dataset, however, is based on the axisymmetric DKI framework and thus might well be confounded by the circularity argument. However, our noise-robustness argument is also a reasonable explanation for the superiority of axisymmetric DKI in this case. Thus, the truth might be in between, i.e., the real improvement of estimation accuracy in the LA dataset when using axisymmetric DKI might be lower than in the simulation but we would expect to still observe an improvement in in-vivo like data.

#### Limits of current measurements protocols

Looking at the estimation accuracy for each of the five AxTM individually revealed that each metric comes with different SNR requirements. Estimation of *W*_∥_ with a MAPE*<* 5%, for example, required an SNR of 81 in the HA voxel of the `synthetic dataset` and an SNR of *>* 100 in the MA voxel of the `synthetic dataset` if RBC was not used. This reveals that current measurement protocols could reach their limits under realistic conditions where the SNR is below 81 or 100 if Rician bias correction is not used. This underlines the importance of using Rician bias correction in cases where all five AxTM are of importance, e.g., for estimation of the biophysical microstructure parameters.

#### Limits of RBC for single voxel application

Similar to previous simulation studies on RBC (Veraart et al., 2011, 2013a; André et al., 2014), we focused on the effects of RBC on the averaged estimated AxTM over the 2500 noise samples. The SNR at which the AxTM could be estimated with a MAPE *<* 5% is reported for that ensemble average. Therefore, the reported results may not directly translate themselves one to one into in-vivo like applications where only one noise realization per voxel is measured.

## 5 Conclusion

Our study revealed that Axisymmetric DKI with RBC is the most SNR effective choice for estimating the AxTM because of two mutually supporting factors. First, RBC itself is most effective for the parallel diffusivity and kurtosis and the mean kurtosis, however, it needs at least some level of fiber alignment to work. Second, compared to standard DKI, axisymmetric DKI is superior in fibers with low alignment and more effective for estimating the perpendicular diffusivity and kurtosis. This makes the combination of axisymmetric DKI with RBC a possibly valuable tool for neuroscience and clinical research studies where a gain in SNR could either be used to reduce scan time or increase spatial resolution.

## Glossary

RBC: Rician bias correction.
DTI: Diffusion tensor imaging.
DKI: Diffusion kurtosis imaging.
HA: Highly aligned fibers.
MA: Fibers with moderate alignment.
LA: Fibers with low alignment.
AxTM: Axisymmetric DKI tensor metrics: *D*_∥_, *D*_⊥_, *W*_∥_, *W*_⊥_, and 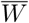.
*D*_∥_: Parallel diffusivity.
*D*_⊥_: Perpendicular diffusivity.
*W*_∥_: Parallel kurtosis.
*W*_⊥_: Perpendicular kurtosis.
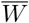: Mean kurtosis.

## 6 Availability of data and materials

The open source ACID toolbox for SPM contains the estimation methods for standard and axisymmetric DKI with and without Rician bias correction used in this study and is available at http://www.diffusiontools.com/.

## 7 Funding

This work was supported by the German Research Foundation (DFG Priority Program 2041 “Computational Connectomics”, [AL 1156/2-1;GE 2967/1-1; MO 2397/5-1; MO 2249/3–1], by the Emmy Noether Stipend: MO 2397/4-1) and by the BMBF (01EW1711A and B) in the framework of ERA-NET NEURON.

## 8 Authors’ contributions

**Jan Malte Oeschger:** Conceptualization, Data curation, Formal analysis, Investigation, Methodology, Software, Visualization, Writing – original draft **Karsten Tabelow:** Conceptualization, Methodology, Supervision, Writing – review & editing, **Siawoosh Mohammadi:** Conceptualization, Funding acquisition, Methodology, Project administration, Resources, Supervision, Writing – review

## Acknowledgements

The implemented code for axisymmetric DKI fitting makes use of the following externally written tools:

– The Gauss Newton fit algorithm implementation used in this study was conceptualized and written by Jan Modersitzki (Modersitzki, 2009) and expanded by Lars Ruthotto who, e.g., implemented slice-wise parameter estimation and introduced an efficient, multi-voxel procedure to accelerate convergence; both improved the algorithm’s run-time.
– For the initial guess of the axisymmetric DKI fit implementation, we used code from the repository of Sune Nørhøj Jespersen: https://github.com/sunenj/Fast-diffusion-kurtosis-imaging-DKI (Hansen et al., 2016).

## Supplementary material

### 8.1 Ground truth DKI datasets

**Table 8.1:** Ground truth in-vivo like standard DKI voxels for the in-vivo like dataset (Figure 2.3), shown are the diffusion and kurtosis tensor components and S_0_, the diffusivities are in 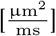.

| Parameter | cb Dataset 1 | cb Dataset 2 | cb Dataset 3 | ct Dataset 1 | ct Dataset 2 | ct Dataset 3 |
| --- | --- | --- | --- | --- | --- | --- |
| $D_{11}$ | 1.92726 | 1.62057 | 1.86790 | 0.53951 | 0.78384 | 0.73087 |
| $D_{22}$ | 0.31583 | 0.38127 | 0.41435 | 0.36193 | 0.42447 | 0.49957 |
| $D_{33}$ | 0.39808 | 0.39813 | 0.36425 | 1.54966 | 1.37428 | 1.29096 |
| $D_{12}$ | -0.00731 | -0.26198 | -0.11110 | -0.15777 | -0.08699 | -0.05191 |
| $D_{13}$ | 0.04079 | -0.25240 | 0.09536 | 0.00094 | -0.08181 | -0.03291 |
| $D_{23}$ | 0.03170 | -0.09149 | -0.03971 | -0.04121 | 0.15430 | 0.03331 |
| $S_0$ | 1.00000 | 1.00000 | 1.00000 | 1.00000 | 1.00000 | 1.00000 |
| $W_{1111}$ | 4.26728 | 4.20207 | 3.73642 | 0.58735 | 1.16398 | 1.12725 |
| $W_{2222}$ | 0.30183 | 0.43656 | 0.36961 | 0.22285 | 0.51330 | 0.74976 |
| $W_{3333}$ | 0.43632 | 0.59204 | 0.26940 | 3.41702 | 2.14949 | 2.30314 |
| $W_{1112}$ | -0.15654 | -0.41618 | -0.18526 | -0.07888 | -0.06150 | -0.08996 |
| $W_{1113}$ | 0.10051 | -0.64422 | 0.18910 | 0.00636 | -0.02166 | 0.02767 |
| $W_{2221}$ | 0.19206 | -0.46340 | -0.26578 | -0.08910 | -0.06061 | -0.09517 |
| $W_{2223}$ | 0.06537 | -0.17548 | -0.08403 | 0.09611 | 0.10937 | -0.04710 |
| $W_{3331}$ | -0.08171 | -0.15145 | 0.12274 | -0.08214 | -0.17008 | -0.09078 |
| $W_{3332}$ | 0.07430 | -0.16995 | -0.09129 | -0.12567 | 0.28125 | -0.09293 |
| $W_{1122}$ | 0.45163 | 0.51896 | 0.50221 | 0.20913 | 0.21167 | 0.25976 |
| $W_{1133}$ | 0.45000 | 0.53712 | 0.43661 | 0.52253 | 0.56751 | 0.30986 |
| $W_{2233}$ | 0.15867 | 0.16724 | 0.07048 | 0.32234 | 0.42040 | 0.39227 |
| $W_{1123}$ | -0.00501 | -0.08834 | -0.03636 | 0.07669 | 0.03705 | -0.00512 |
| $W_{2213}$ | -0.00262 | 0.00521 | 0.08195 | -0.02098 | -0.08206 | -0.02479 |
| $W_{3312}$ | 0.03732 | -0.13410 | -0.11029 | -0.16710 | -0.06077 | -0.02733 |
| Parameter | or Dataset 1 | or Dataset 2 | or Dataset 3 | slf Dataset 1 | slf Dataset 2 | slf Dataset 3 |
| $D_{11}$ | 1.06085 | 0.67273 | 0.69468 | 0.46699 | 0.68565 | 0.63911 |
| $D_{22}$ | 0.59047 | 0.75000 | 1.79639 | 0.43803 | 0.48026 | 0.61127 |
| $D_{33}$ | 1.36088 | 1.48673 | 0.49756 | 1.69614 | 1.23269 | 1.22297 |
| $D_{12}$ | 0.12149 | 0.04143 | -0.43159 | 0.05648 | 0.02784 | 0.08187 |
| $D_{13}$ | 0.61736 | 0.22247 | 0.19896 | 0.07981 | -0.07817 | -0.04450 |
| $D_{23}$ | 0.05184 | 0.26023 | -0.21783 | -0.20355 | -0.15512 | -0.09252 |
| $S_0$ | 1.00000 | 1.00000 | 1.00000 | 1.00000 | 1.00000 | 1.00000 |
| $W_{1111}$ | 1.18421 | 0.74417 | 0.60364 | 0.38485 | 1.27168 | 0.65976 |
| $W_{2222}$ | 0.48582 | 0.71486 | 2.21601 | 0.51498 | 0.72106 | 0.83091 |
| $W_{3333}$ | 1.72750 | 1.99702 | 0.37346 | 3.25042 | 2.56281 | 1.96113 |
| $W_{1112}$ | 0.09557 | 0.04945 | -0.18369 | 0.03519 | 0.11477 | 0.00824 |
| $W_{1113}$ | 0.42320 | 0.11965 | 0.04012 | 0.01711 | 0.05484 | 0.06571 |
| $W_{2221}$ | 0.22072 | -0.01878 | -0.49678 | 0.11049 | -0.03851 | -0.10122 |
| $W_{2223}$ | 0.13593 | -0.00500 | -0.40378 | -0.02510 | 0.06858 | 0.06213 |
| $W_{3331}$ | 0.70421 | 0.13580 | 0.03943 | 0.15730 | -0.32397 | -0.13608 |
| $W_{3332}$ | 0.15332 | 0.36637 | 0.09475 | -0.41687 | -0.24668 | -0.29428 |
| $W_{1122}$ | 0.19154 | 0.25614 | 0.52836 | 0.12737 | 0.22065 | 0.18565 |
| $W_{1133}$ | 0.61697 | 0.26980 | 0.15419 | 0.49145 | 0.36651 | 0.53366 |
| $W_{2233}$ | 0.26455 | 0.40726 | 0.34763 | 0.42856 | 0.14162 | 0.27141 |
| $W_{1123}$ | 0.04012 | 0.00642 | -0.11345 | -0.04016 | -0.01867 | -0.08256 |
| $W_{2213}$ | 0.06872 | 0.07859 | 0.24902 | -0.01085 | -0.04099 | 0.02417 |
| $W_{3312}$ | 0.02283 | 0.04014 | -0.03715 | 0.06156 | 0.07493 | 0.17797 |

**Table 8.2:**
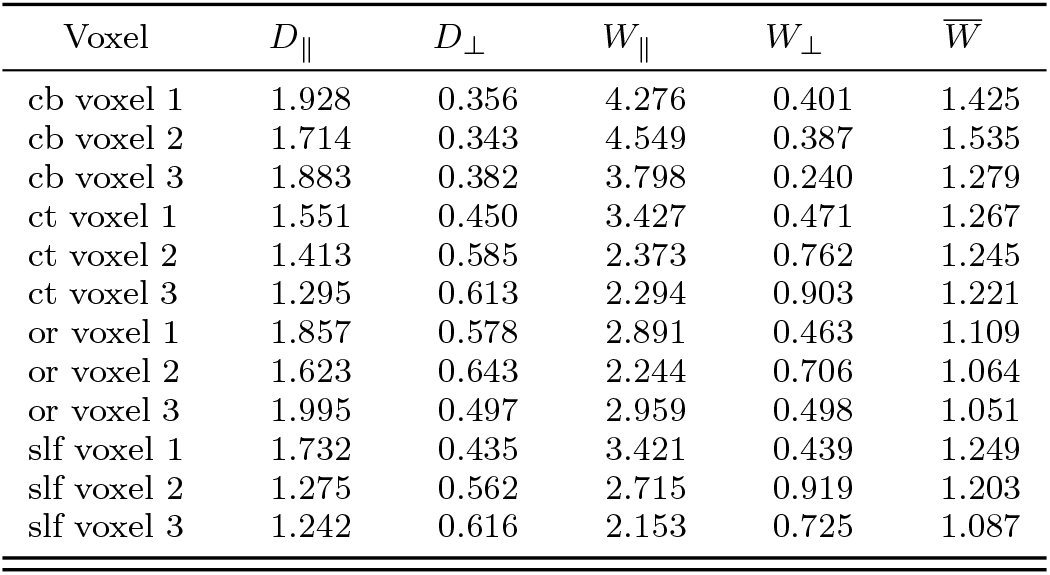
Ground truth AxTM of the in-vivo like dataset, corresponding to the tensor components listed in Table 8.1, the diffusivities are in 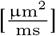.

| Voxel | $D_{\parallel}$ | $D_{\perp}$ | $W_{\parallel}$ | $W_{\perp}$ | $\bar{W}$ |
| --- | --- | --- | --- | --- | --- |
| cb voxel 1 | 1.928 | 0.356 | 4.276 | 0.401 | 1.425 |
| cb voxel 2 | 1.714 | 0.343 | 4.549 | 0.387 | 1.535 |
| cb voxel 3 | 1.883 | 0.382 | 3.798 | 0.240 | 1.279 |
| ct voxel 1 | 1.551 | 0.450 | 3.427 | 0.471 | 1.267 |
| ct voxel 2 | 1.413 | 0.585 | 2.373 | 0.762 | 1.245 |
| ct voxel 3 | 1.295 | 0.613 | 2.294 | 0.903 | 1.221 |
| or voxel 1 | 1.857 | 0.578 | 2.891 | 0.463 | 1.109 |
| or voxel 2 | 1.623 | 0.643 | 2.244 | 0.706 | 1.064 |
| or voxel 3 | 1.995 | 0.497 | 2.959 | 0.498 | 1.051 |
| slf voxel 1 | 1.732 | 0.435 | 3.421 | 0.439 | 1.249 |
| slf voxel 2 | 1.275 | 0.562 | 2.715 | 0.919 | 1.203 |
| slf voxel 3 | 1.242 | 0.616 | 2.153 | 0.725 | 1.087 |

**Table 8.3:** Set of synthetic AxTM, *S*_0_ and axis of symmetry 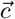 used to simulate the synthetic dataset based on axisymmetric DKI. The synthetic dataset consisting of three voxels with sets of 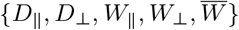 was taken from Coelho et al. (2019), diffusivities are in 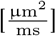, S is in arbitrary units.

| Dataset | $D_{\parallel}$ | $D_{\perp}$ | $W_{\parallel}$ | $W_{\perp}$ | $\bar{W}$ | $S_0$ | $\vec{c}$ |
| --- | --- | --- | --- | --- | --- | --- | --- |
| Fibers with high alignment (HA) | 1.503 | 0.195 | 1.456 | 0.291 | 0.926 | 1 | $(1, 0, 0)^T$ |
| Fibers with moderate alignment (MA) | 1.557 | 1.048 | 0.396 | 0.708 | 0.330 | 1 | $(1, 0, 0)^T$ |
| Fibers with low alignment (LA) | 0.457 | 0.408 | 2.901 | 2.702 | 2.770 | 1 | $(1, 0, 0)^T$ |

## Notes

https://bitbucket.org/siawoosh/acid-artefact-correction-in-diffusion-mri/src/master/

### Competing Interest Statement

The authors have declared no competing interest.

